# Mistranslating tRNA identifies a deleterious S213P mutation in the *Saccharomyces cerevisiae eco1-1* allele

**DOI:** 10.1101/2020.04.08.031922

**Authors:** Yanrui Zhu, Matthew D. Berg, Phoebe Yang, Raphaël Loll-Krippleber, Grant W. Brown, Christopher J. Brandl

## Abstract

Mistranslation occurs when an amino acid not specified by the standard genetic code is incorporated during translation. Since the ribosome does not read the amino acid, tRNA variants aminoacylated with a non-cognate amino acid or containing a non-cognate anticodon dramatically increase the frequency of mistranslation. In a systematic genetic analysis, we identified a suppression interaction between tRNA^Ser^UGG, G26A, which mistranslates proline codons by inserting serine, and *eco1-1,* a temperature sensitive allele of the gene encoding an acetyltransferase required for sister chromatid cohesion. The suppression was partial with a tRNA that inserts alanine at proline codons and not apparent for a tRNA that inserts serine at arginine codons. Sequencing of the *eco1-1* allele revealed a mutation that would convert the highly conserved serine 213 within β7 of the GCN5-related N-acetyltransferase core to proline. Mutation of P213 in *eco1-1* back to the wild-type serine restored function of the enzyme at elevated temperature. Our results indicate the utility of mistranslating tRNA variants to identify functionally relevant mutations and identify *eco1* as a reporter for mistranslation. We propose that mistranslation could be used as a tool to treat genetic disease.

## INTRODUCTION

Mistranslation occurs when an amino acid that differs from that specified by the standard genetic code is incorporated into nascent proteins during translation. Mistranslation naturally occurs at a frequency of approximately one in ten thousand codons in all cells and increases under specific environmental conditions or upon mutation of the translational machinery (Santos *et al.* 1999; Bacher *et al.* 2007; Ling *et al.* 2007; Kramer and Farabaugh 2007; Drummond and Wilke 2009; Javid *et al.* 2014). Contrary to Crick’s Frozen Accident Theory (Crick 1968), mistranslation is tolerated at levels approaching 8% (Berg *et al.* 2019). Accurate translation has two major components. The first is tRNA aminoacylation, catalyzed by aminoacyl-tRNA synthetases (aaRS) that specifically couple an amino acid to the 3’ end of their cognate tRNA(s) [reviewed in Pang *et al.* (2014)]. The second specificity step is codon decoding at the ribosome, which relies on base pairing between codon and anticodon. Loss of fidelity at either step can lead to mistranslation. Mistranslation is dramatically increased by tRNA variants that are inaccurately aminoacylated or contain mutations within their anticodon (Geslain *et al.* 2010; Hoffman *et al.* 2017; Lant *et al.* 2017; Berg *et al.* 2017; Zimmerman *et al.* 2018). Nucleotides in the tRNA that are recognized by a specific aaRS are called *identity elements* and consist of single nucleotides, nucleotide pairs, and structural motifs (de Duve 1988; Hou and Schimmel 1988; Giegé *et al.* 1998). Since the anticodon links an amino acid to its codon assignment, it is not surprising that tRNA recognition often involves identity elements within the anticodon. However, for tRNA^Ser^ and tRNA^Ala^ the anticodon plays no role in the specificity of charging in yeast and for tRNA^Leu^ the anticodon only plays a minor role (Giegé *et al.* 1998), making these tRNAs particularly amenable to engineering for increased mistranslation.

Studies demonstrating the ability of tRNA variants to correct genetic errors by replacing a non-functional residue with a functionally competent residue predate the deciphering of the genetic code (Crawford and Yanofsky 1959; Stadler and Yanofsky 1959; Yanofsky and Crawford 1959). Intergenic suppressors that change codon meaning were called informational suppressors, since they alter the information flow from DNA to protein (Gorini and Beckwith 1966). Some of the early studies included a demonstration by Benzer and Champe (1962) of the suppression of nonsense mutations. They reasoned that suppression acts by changing the genetic code to add a new sense codon. Mutations in the *Escherichia coli* tryptophan synthase (trpA) gene provided a selection to identify mistranslation inducing mutations that rescued growth in tryptophan free media. These studies characterize tRNA variants that led to Gly insertion at Arg or Cys codons (Carbon *et al.* 1966; Jones *et al.* 1966; Gupta and Khorana 1966). Other suppressor mistranslating tRNAs have been identified in yeast. For example, Goodman *et al.* (1977) mapped a tRNA^Tyr^ with a G to T transversion mutation resulting in a U at the wobble nucleotide position making it a nonsense suppressor.

Previously, we engineered serine tRNA variants that mis-incorporate serine at proline codons by replacing the UGA anticodon with UGG (Berg *et al.* 2017, 2019). These tRNAs contain secondary mutations to reduce tRNA functionality and modulate mistranslation levels since plasmids expressing a tRNA with this anticodon change alone can not be transformed into yeast. One variant, tRNA^Ser^_UGG, G26A_, contains a G26A secondary mutation and when expressed from a centromeric plasmid results in a frequency of serine incorporation at proline codons of ~ 5% as determined by mass spectrometry (Berg *et al.* 2019). Zimmerman *et al.* (2018) and Geslain *et al.* (2010) have also demonstrated the possibility of mistranslating serine for a number of amino acids in yeast and mammalian cells.

In this report we use a mistranslating tRNA variant to identify the causative mutation in the *Saccharomyces cerevisiae eco1-1* allele as a serine to proline missense mutation. Eco1 is an acetyltransferase required for sister chromatid cohesion during DNA replication (Tóth *et al.* 1999; Unal *et al.* 2007; Ben-shahar *et al.* 2008). Mutations in the human homolog of *ECO1* (ESCO2) cause Roberts syndrome (Vega *et al.* 2005), a rare genetic disorder characterized by limb reduction and craniofacial abnormalities. We demonstrate the utility of mistranslation as a tool to identify causative mutations, demonstrate *eco1* as a selectable reporter to monitor mistranslation and provide support for the possibility of using mistranslation as a tool to cure genetic disease.

## MATERIALS AND METHODS

### Yeast strains and DNA constructs

The SGA starter strain, Y7092 *(MATα can1Δ::STE2pr-SpHIS5 lyp1Δ his3Δ1 leu2Δ0 ura3Δ0 met15Δ0*), was a kind gift from Dr. Brenda Andrews (University of Toronto). Strains from the temperature sensitive collection are derived from the wild-type *MAT**a*** haploid yeast strain BY4741 (*MAT***a** *his3Δ0 leu2Δ0 met15Δ0 ura3Δ0;* Winzeler and Davis 1997) and described in Costanzo *et al.* (2016). CY8613 (*MAT*α *HO::natNT2-SUP17_UGG, G26A_ can1Δ::STE2pr-SpHIS5 lyp1Δ)* contains a gene encoding tRNA^Ser^_UGG, G26A_ integrated into Y7092 at the *HO* locus as described below. The isogenic control strain lacking the mistranslating tRNA encoding gene is CY8611 (*MAT*α *HO::natNT2 can1Δ::STE2pr-SpHIS5 lyp1Δ*).

Yeast strains were grown in yeast peptone media containing 2% glucose (YPD) or synthetic media supplemented with nitrogenous bases and amino acids (SD) at the temperature indicated. Plates lacking uracil (-URA) were made in SD media supplemented with 0.6% (g/vol) casamino acids, 0.25% adenine and 0.5% tryptophan. The temperature sensitive collection was maintained in 1536-format on YPD plates containing 200 mg/L geneticin (G418; Invitrogen). The SGA query strain was maintained on YPD plates containing 100 mg/L nourseothricin-dihydrogen sulfate (NAT; Werner BioAgents). Double mutants containing both the SGA query and temperature sensitive mutation were maintained on synthetic dropout plates lacking histidine, arginine and lysine with monosodium glutamate (1 g/L) as the nitrogen source, canavanine (50 mg/L), thialysine (50 mg/L), G418 (200 mg/L) and NAT (100 mg/L).

Centromeric plasmids expressing tRNA^Ser^ (pCB3076), tRNA^Ser^_UGG, G26A_ (pCB4023), tRNA^Pro^_UGG G3:U70_ (pCB2948) and tRNA^Ser^_UCU, G26A_ (pCB4301) are described in Berg *et al.* (2017), Hoffman *et al.* (2017) and Berg *et al.* (2019).

### Construction of SGA Query and Control Strains

The protocol for constructing the SGA query strains was adapted from the PCR-mediated gene deletion method of Tong *et al.* (2001). A DNA fragment containing 200 bp of upstream *HO* flanking region and 200 bp of the *HO* gene was synthesized by Life Technologies and cloned into pGEM^®^-T Easy (Promega Corp.) as a *NotI* fragment (pCB4386). The *natNT2* marker from pFA6a-natNT2 was PCR amplified using primers UK9789/UK9790 (Table 1) and cloned into pCB4386 as an *Eco*RI fragment to generate the control SGA integrating vector (pCB4394). The gene encoding tRNA^Ser^_UGG, G26A_ was PCR amplified from pCB4023 (Berg *et al.* 2017) using primers UG5953/VB2609 and inserted as a *HindIII* fragment into pCB4394 to generate pCB4397. pCB4394 and pCB4397 were digested with *Not*I, transformed into Y7092 and transformants selected on 100 mg/L NAT to generate the SGA strains CY8611 and CY8613, respectively. Integration of the fragments was verified by PCR.

**Table 1.**
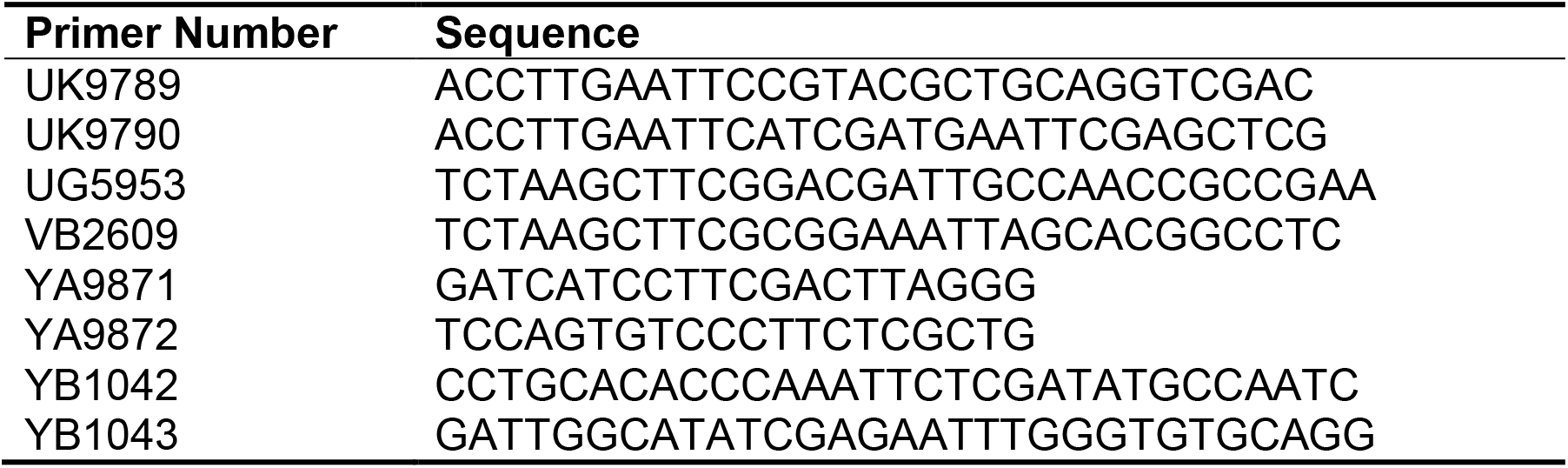
Primers used in this study

### Synthetic genetic array analysis and validation

The SGA assay was performed as described by Tong *et al.* (2001). The SGA control and query strains (CY8611 and CY8613) were mated to a temperature-sensitive collection (Ben-Aroya *et al.* 2008; Li *et al.* 2011; Kofoed *et al.* 2015; Costanzo *et al.* 2016) arrayed in quadruplicate 1536-array format on YPD plates using a BioMatrix robot (S&P Robotics Inc.). Mated strains were grown overnight then pinned onto YPD + NAT/G418 plates to select for diploids. Haploids were generated by pinning the diploid strains onto enriched sporulation plates and incubating for 1 week at 22°C. The haploids then underwent three rounds of selection for double mutants that had both the tRNA mutation and temperature-sensitive allele. First, strains were pinned on SD – His/Arg/Lys + canavanine/thialysine plates to select for MATa haploids. Next, colonies were pinned twice onto (SD/MSG) - His/Arg/Lys + canavanine/thialysine/G418/NAT to select double mutants. Colonies were incubated for two days between pinnings at room temperature. The double mutants were grown at 30°C for 5 days. Images from day 3 were analyzed and scored using SGAtools (Wagih *et al.* 2013). An SGA score ≥ 0.5 and a p-value ≤ 0.05 was used to identify alleles that were potentially suppressed by tRNA^Pro^_UGG, G26A_. Suppression was validated by transforming *URA3* centromeric plasmids expressing tRNA^Pro^_UGG, G26A_ (pCB4023) or wild-type tRNA^Ser^ (pCB3076; Berg *et al.* 2017) into each strain and comparing growth on plates lacking uracil.

### Isolation of *eco1-1* and mutagenesis

Genomic DNA was isolated from the temperature sensitive *eco1-1* strain as described by Hoffman and Winston (1987). *eco1-1* was PCR amplified using primers YA9871/YA9872. The PCR included Q5^®^ High-Fidelity DNA Polymerase (New England Biolabs) to minimize incorporation errors. The PCR product was sequenced with primer YA9872 and cloned into pGEM^®^-T Easy (pCB4639) where it was re-sequenced with M13 forward and reverse primers. Pro213 was mutagenized to Ser213 by two-step PCR mutagenesis. Primer pairs YA9871/YB1043 and YA9872/YB1042 were used in the first round followed by outside primers YA9871 and YA9872. The PCR product was subcloned into pGEM^®^-T Easy and cloned as an *Eco*RI fragment into YCplac33 to give pCB4673. Similarly, *eco1-1* was subcloned into YCplac33 to give pCB4662.

### Modeling

The protein sequence for *Saccharomyces cerevisiae* Eco1 (SGD ID: S000001923) was aligned to *Homo sapiens* ESCO1 (Uniprot: Q5FWF5-1) and ESCO2 (Uniprot: Q56NI9-1), *Mus musculus* Esco1 (Uniprot: Q69Z69) and Esco2 (Uniprot: Q8CIB9), *Danio rerio* esco1 (Uniprot: X1WEK0) and esco2 (Uniprot: Q5SPR8) and *Drosophila melanogaster* eco (Uniprot: Q9VS50) using Clustal Omega (Madeira *et al.* 2019). The S213P mutation, corresponding to S770, was modelled on the human ESCO1 structure (PDB: 4MXE; Kouznetsova *et al.* 2016) using Missense3D (Ittisoponpisan *et al.* 2019).

## RESULTS

We performed an SGA screen to identify genes demonstrating genetic interactions with tRNA^Ser^_UGG, G26A_, a tRNA variant that mistranslates serine at proline codons, using a temperature sensitive collection containing 1016 temperature sensitive alleles (Kofoed *et al.* 2015; Costanzo *et al.* 2016). The tRNA encoding gene was integrated at the *HO*analysis demonstrates the utility of mistranslation to locus and selected for by NAT resistance. The control strain contained the *natNT2* marker integrated at *HO*, but no tRNA. Screens were performed at 30°C and analyzed using SGAtools (Wagih *et al.* 2013). The negative genetic interactions identified will be described elsewhere. In this screen, the *eco1-1* strain had a genetic interaction score of 0.52 *(P* = 1.0 x 10^-5^) suggesting it grew better than expected in the presence of tRNA^Ser^_UGG, G26A_. To validate the positive genetic interaction, centromeric plasmids carrying the gene encoding tRNA^Ser^_UGG, G26A_ (pCB4023) or wild-type tRNA^Ser^ (pCB3076) were transformed into the temperature sensitive strain and its growth compared on plates lacking uracil at 24°C, 30°C and 37°C. As shown in Figure 1A, tRNA^Ser^_UGG, G26A_ improved growth of the *eco1-1* strain at 30°C and 37°C. Note that the partial toxicity of the mistranslating tRNA (tRNA^Ser^_UGG, G26A_) is apparent when the cells are grown at 24°C.

**Figure 1.**
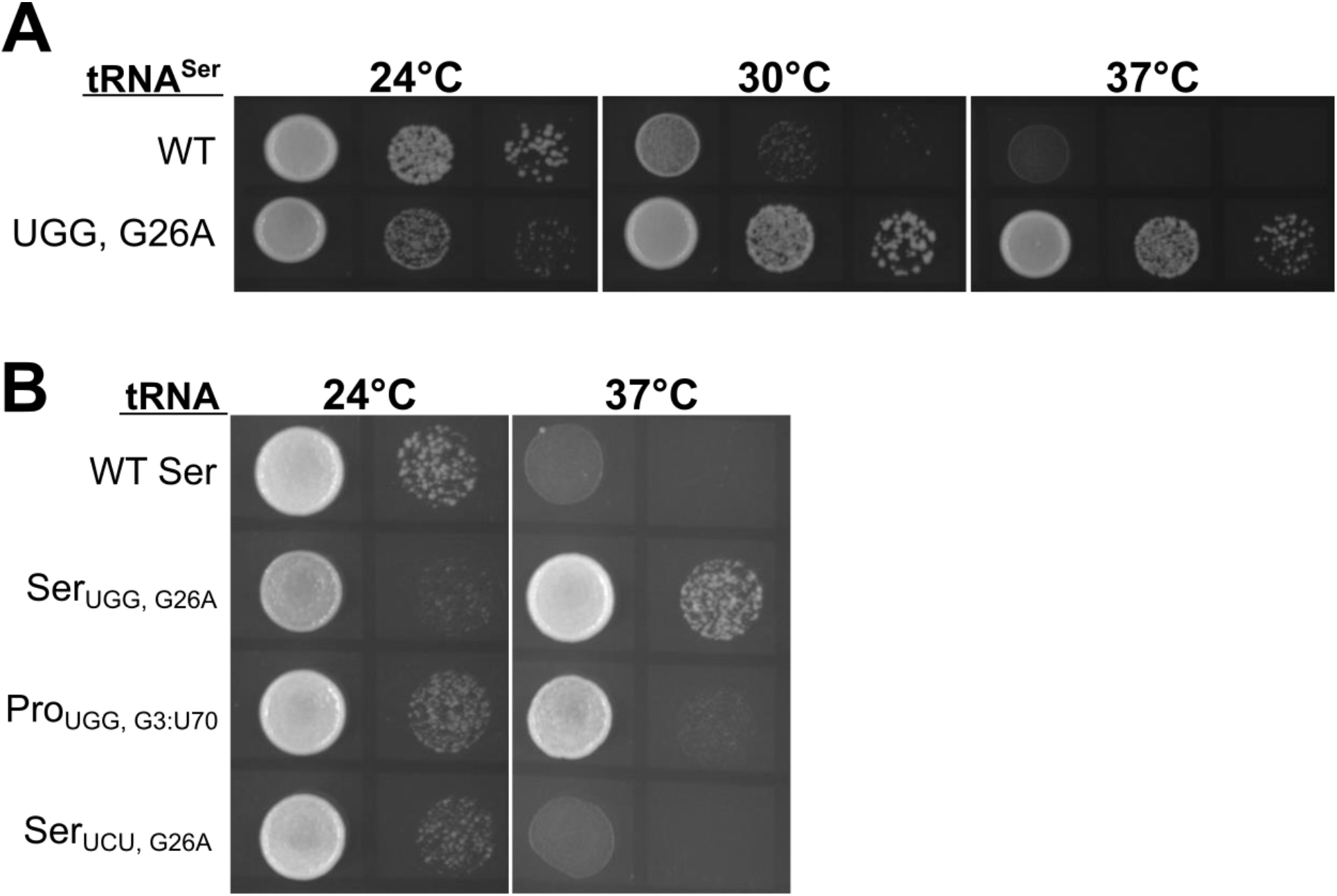
Suppression of a temperature sensitive eco1-1 allele through mistranslation. **A.** The *eco1-1* yeast strain was transformed with *URA3* centromeric plasmids expressing tRNA^Ser^ (pCB3076) or tRNA^Ser^_UGG, G26A_ (pCB4023), grown to saturation in - URA medium at room temperature, spotted in 10-fold serial dilutions on -URA plates and grown at 24°C, 30°C or 37°C. Images were taken after two days of growth. **B.** The *eco1-1* strain from the temperature sensitive collection was transformed with centromeric plasmids expressing tRNA^Ser^ (pCB3076), tRNA^Ser^_UGG, G26A_ (pCB4023), tRNA^Pro^_UGG, G3:U70_ (pCB2948) or tRNA^Ser^_UCU, G26A_ (pCB4301). Cells were grown to saturation in -URA medium, spotted in 10-fold serial dilutions on -URA plates and grown for two days at 24°C or 37°C.

The suppression could arise from a mistranslation event, effectively reverting the mutation or from a positive interaction between *eco1-1* and mistranslation in general. To distinguish between these possibilities, we transformed the *eco1-1* strain with plasmids expressing tRNA^Ser^, tRNA^Ser^_UGG, G26A_, tRNA^Pro^_G3:U70_ (which inserts alanine at proline codons) and tRNA^Ser^_UCU, G26A_ (which inserts serine at arginine codons) (Figure 1B). Partial suppression was seen with tRNA^Pro^_G3:U70_. No suppression was seen with tRNA^Ser^_UCU, G26A_ or with wild-type tRNA^Ser^. We conclude that the tRNA suppresses the allele by mistranslation at a proline codon, rather than through a more general genetic interaction between *eco1-1* allele and mistranslation and that misincorporation of serine at proline codons results in stronger suppression of *eco1-1* than does misincorporation of alanine.

Based on the suppression by tRNA^Ser^_UGG, G26A_ (substitutes serine at proline codons) and to a lesser extent tRNA^Pro^_UGG, G3:U70_ (substitutes alanine at proline codons) we predicted that the *eco1-1* strain in the collection contains a mutation resulting in the conversion of a serine residue to proline that was contributing to the temperature sensitive phenotype. We isolated the *eco1-1* gene, including up- and down-stream flanking sequence by PCR and cloned the product. The clone was sequenced through the gene. We identified four missense mutations, G184D, S213P, K260R and G273D, and one synonymous mutation (Table 2). The missense mutation altering S213 to proline converts the UCG codon for serine to CCG for proline. We have previously shown that CCG is mistranslated to serine by tRNA^Ser^_UGG, G26A_ (Berg *et al.* 2019). As an indication that the serine to proline mutation was not a clonal artifact, the PCR product was directly sequenced with the 3’ oligonucleotide, which confirmed the presence of the T to C mutation at nucleotide 637 of *eco1-1.* We note that Tóth *et al.* (1999) characterized a temperature sensitive allele of *eco1* with a mutation converting glycine 211 to aspartic acid that they named *eco1-1.* It is questionable whether this is the allele in the collection since their study was performed with a W303 strain background.

**Table 2.**
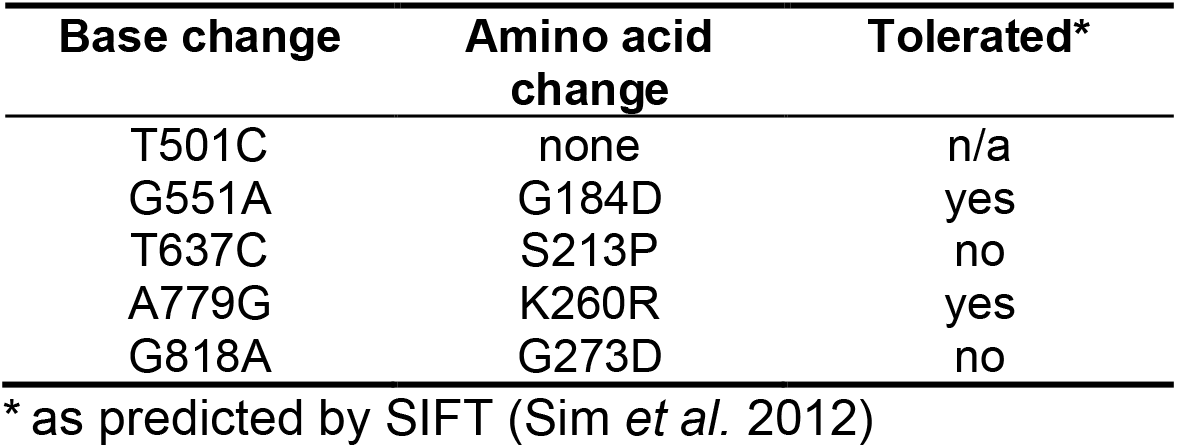
Mutations in *eco1-1*

To analyze which of the four missense mutations (G184D, S213P, K260R or G273D) resulted in changes to important regions of the protein we performed an alignment of Eco1 homologs from yeast, human, mouse, fruit fly and zebrafish. Of these only S213 is in a highly conserved region of the protein (Figure 2A). Furthermore, analysis by SIFT (Sim *et al.* 2012) suggests that any change of S213 would be detrimental to function. Eco1 encodes a histone acetyltransferase required for chromatid cohesion during DNA replication. Mutations in the human gene (ESCO2) cause Roberts syndrome (Vega *et al.* 2005). The structure of ESCO2 has been determined (Kouznetsova *et al.* 2016; Rivera-Colón *et al.* 2016). S770, the equivalent of yeast S213, is found in β7 strand of the GCN5-related N-acetyltransferase core (Figure 2B). Though not essential for function, mutation of S770 (yeast S213) to alanine reduces catalytic efficiency ~8-fold *in vitro,* with a minimal effect on thermal stability (Rivera-Colón *et al.* 2016). Modeling of a proline at this position 213 suggests that this would distort the β7 strand perhaps altering the stability of the protein (Figure 2C).

**Figure 2.**
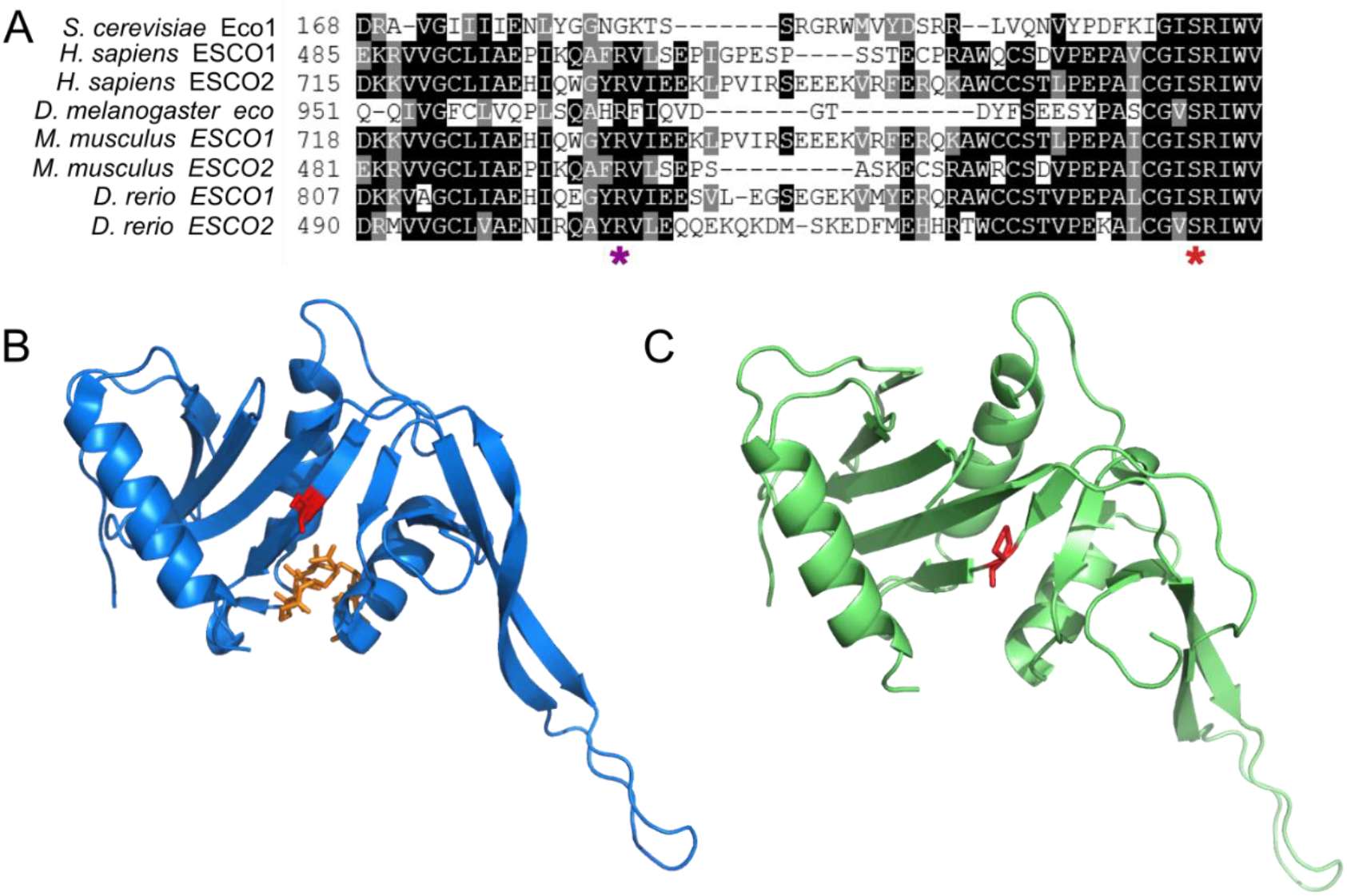
Sequence alignments and modelling of eco1-1 suggest S213P could disrupt function. **A.** *S. cerevisiae* Eco1 was aligned to homologs from *Homo sapiens, Drosophila melanogaster*, *Mus musculus* and *Danio rerio* using Clustal Omega (Madeira *et al.* 2019). The purple star indicates the G184 which was found to be mutated to D in *eco1-1* and the red star indicates S213 found to be mutated to proline. **B.** Residue S770 in human ESCO1, corresponding to yeast S213, is highlighted in red on the human ESCO1 structure [4MXE; Kouznetsova *et al.* (2016)]. The acetyl-CoA substrate is shown in orange. **C.** Missense3D (Ittisoponpisan *et al.* 2019) was used to model the yeast S213P mutation onto the human ESCO1 structure. The red residue indicates S770P.

To determine if the S213P mutation resulted in the temperature sensitive nature of *eco1-1*, we constructed the allele where only P213 was converted back to TCG for serine. *eco1-1* and *eco1-1_P213S_* alleles were inserted into a *URA3* containing centromeric plasmid and transformed into the *eco1-1* strain. The ability of this *eco1-1_P213S_* allele, the original *eco1-1* allele (carrying four mutations) and the plasmid alone to complement the temperature sensitive nature of the *eco1-1* strain was tested by analyzing growth at 24°C and 37°C (Figure 3). At 24°C the *eco1-1* strain grows well with or without an additional copy of the *eco1* allele. At 37°C an additional *eco1-1* allele (with P213) allowed for slightly better growth than empty plasmid (Figure 3). In contrast addition of the wild-type-like S213 *eco1-1* allele *(eco1-1_P213S_)* restored robust growth of the strain at 37°C. This result confirms that mutation of wild-type serine at 213 to proline is the cause of the temperature sensitive nature of the *eco1-1* allele and that mistranslation of the proline codon in *eco1-1* to insert serine would be sufficient to allow growth at elevated temperature.

**Figure 3.**
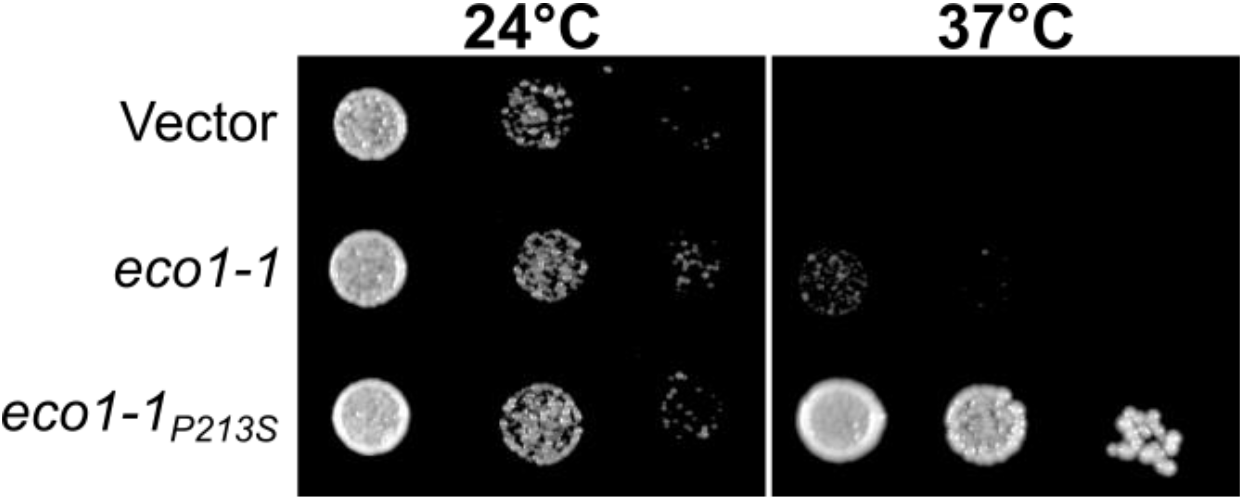
Reversion of P213 in eco1-1 back to the wild-type serine allows robust growth at elevated temperatures. The *eco1-1* yeast strain was transformed with either an empty *URA3* centromeric plasmid or a *URA3* plasmid expressing *eco1-1* (pCB4639) or *eco1-1_P213S_* (pCB4673), grown to saturation in -URA medium at room temperature, spotted in 10-fold serial dilutions on -URA plates and grown at 24°C or 37°C. Images were taken after two days of growth.

## DISCUSSION

Our analysis demonstrates the utility of mistranslation to identify the nature of deleterious mutations. Suppression of the *eco1-1* temperature sensitive phenotype with tRNA^Ser^_UGG, G26A_, which causes serine mistranslation at proline codons (Berg *et al.* 2017), and to a lesser extent tRNA^Pro^_G3:U70_, which causes alanine mistranslation at proline codons (Hoffman *et al.* 2017), suggested that a missense mutation creating a proline codon was involved. Upon sequencing of the *eco1-1* allele from the temperature sensitive collection, we found a missense mutation converting S213 to proline. This mutation is in a highly conserved region of the acetyltransferase domain. The contribution of the S213P mutation to the temperature sensitivity of *eco1-1* was demonstrated by converting it back to a serine codon. In the context of the other mutations in the *eco1-1* allele, the reversion was sufficient to complement the temperature sensitive phenotype.

Aminoacylation of the tRNAs for serine and alanine and to a lesser extent leucine do not depend on the anticodon in yeast. This allows the construction of tRNAs that insert these amino acids at non-cognate codons. Many of these tRNAs have been engineered (for example see Geslain *et al.* 2010; Zimmerman *et al.* 2018; Berg *et al.* 2019). Through analysis similar to that performed here, these mistranslating tRNAs will allow the identification of functionally significant missense mutations in alanine, serine and leucine codons. The one caveat to the method is that mistranslation frequency has a threshold of approximately 8% in yeast due to introducing proteotoxic stress. The protein in question must therefore show significant function when present at relatively low levels. Other examples of proteins that function at such reduced levels are the proline isomerase Ess1 (Gemmill *et al.* 2005) and the cochaperone Tti2 (Hoffman *et al.* 2016, 2017).

If one assumes that all mutations have equally likelihood of creating a temperature sensitive allele, the maximum number of temperature sensitive strains in a collection that contain a causative serine to proline mutation can be estimated from the frequency of UCA (0.89%) and UCG (0.85%) serine codons. With a single base change, each of these codons becomes efficiently decoded by the UGG anticodon. The combined usage frequency of UCA and UCG predicts that ~18 of the 1016 strains, contain a serine to proline mutation. As we identified one strain suppressed by tRNA^Ser^_UGG, G26A_ in the collection, it suggests that for ~ 6% (1/18) of essential yeast proteins, 5% of their native expression (the extent of proline to serine mistranslation) is sufficient to support viability.

Functional assays that result in phenotypic reversion are one of the simplest methods to screen for mistranslation events. We previously used a leucine to proline mutation in *TTI2* to identify mistranslating tRNA variants (Hoffman *et al.* 2017; Berg *et al.* 2017). The utility of using *tti2* to detect different varieties of mistranslation caused by tRNA variants is somewhat limited because our screening has only revealed leucine to proline mutations to have phenotypic consequences when found in isolation. *eco1-1* may provide a more versatile reporter in that the structure is known and altering the acetyltransferase domain would be expected to impact structure or function in a way that results in phenotypic change. We note in particular the conservation of the residues flanking S213 (Figure 2A), making those codons candidates for further reporter engineering.

Many diseases result from missense or nonsense mutations that alter the structure, function and/or stability of a gene product. Our findings complement existing reports documenting the ability of tRNA variants to restore protein function in yeast and bacteria through their ability to mistranslate a deleterious codon (for examples, see reviews by Celis and Piper 1981 and Murgola 1985). Effectively, phenotypic suppression by mutant tRNAs is evidence that mistranslation is able to cure genetic disease. The mistranslation could be achieved through tRNA variants, reducing specificity of aminoacyl-tRNA-synthetases, or decreasing proofreading functions. In contrast to the concerns addressed by Crick in his “Frozen Accident Theory” (Crick 1968), mistranslation is not catastrophic for cell viability. At levels in the 3-5% range mistranslation has minimal affect on yeast cell viability (Hoffman *et al.* 2017; Lant *et al.* 2017; Berg *et al.* 2019). For some key cellular proteins, 3-5% mistranslation is sufficient to restore viability.

## ACKNOWLEDGMENTS

We thank Julie Genereaux for technical assistance and for comments on the manuscript.

